# The diallelic self-incompatibility system in Oleaceae is controlled by a hemizygous genomic region expressing a gibberellin pathway gene

**DOI:** 10.1101/2023.12.18.571976

**Authors:** Vincent Castric, Rita A. Batista, Amélie Carré, Soraya Mousavi, Clément Mazoyer, Cécile Godé, Sophie Gallina, Chloé Ponitzki, Anthony Theron, Arnaud Bellec, William Marande, Sylvain Santoni, Roberto Mariotti, Andrea Rubini, Sylvain Legrand, Sylvain Billiard, Xavier Vekemans, Philippe Vernet, Pierre Saumitou-Laprade

## Abstract

Sexual reproduction in flowering plants is commonly controlled by self-incompatibility (SI) systems that are either homomorphic (and typically governed by large numbers of distinct allelic specificities), or heteromorphic (and then typically governed by only two allelic specificities). The SI system of the Oleaceae family is a striking exception to this rule and represents an evolutionary conundrum, with the long-term maintenance of only two allelic specificities, but often in the complete absence of morphological differentiation between them. To elucidate the genomic architecture and molecular bases of this highly unusual SI system, we obtained chromosome-scale genome assemblies of *Phillyrea angustifolia* individuals belonging to the two SI specificities and connected them to a genetic map. Comparison of the S-locus region revealed a segregating 543-kb indel specific to one of the two specificities, suggesting a hemizygous genetic architecture. Only one of the predicted genes in this indel is conserved with the olive tree *Olea europaea,* where we also confirmed the existence of a segregating hemizygous indel. We demonstrated full association between presence/absence of this gene and the SI groups phenotypically assessed across six distantly related Oleaceae species. This gene is predicted to be involved in catabolism of the Gibberellic Acid (GA) hormone, and experimental manipulation of GA levels in developing buds modified the male and female SI responses in an S-allele-specific manner. Thus, our results provide a unique example of a reproductive system where a single conserved gibberellin-related gene in a 500-700kb hemizygous indel underlies the long-term maintenance of two groups of reproductive compatibility.

## INTRODUCTION

In most eukaryotic species, sexual reproduction is only possible between mating partners belonging to a discrete number of reproductive groups (Billiard et al. 2011). Understanding how these reproductive groups are genetically determined and the long-term evolutionary consequences of their existence are central issues in evolutionary biology, with major implications on biological processes as diverse as the evolution of anisogamy (Ferris et al. 2010), of uniparental inheritance of cytoplasmic DNA (Hadjivasiliou et al 2013) or of sex- chromosomes (Hartmann et al. 2021). In the most familiar case, the reproductive groups correspond to two separate sexes (males and females) that differ by conspicuous phenotypic traits beyond their sheer reproductive compatibility. However this does not need to be the case, and in many situations the reproductive groups can be indistinguishable at the morphological level. They are then more commonly referred to as “mating types” and are controlled by a variety of molecular mechanisms, as documented *e.g.* in fungi, yeasts or green algae.

In hermaphroditic flowering plants, patterns of reproductive compatibility are commonly governed by genetic self-incompatibility (SI) systems (Takayama & Isogai, 2005). Such systems occur in about 40% of flowering plants (Igic et al. 2008) and are thought to have independently evolved multiple times, generating a wide diversity of SI systems. In spite of decades of effort, the genetic architecture and molecular mechanisms underlying SI have only been deciphered in a handful of plant families among the myriads known to exhibit SI. Notwithstanding this, two largely distinct types of genetic architectures and molecular mechanisms of SI have been described (de Nettancourt 1977): i) heteromorphic SI systems, which typically show two (sometimes three) categories of mating partners, with distinct flower morphologies encoded by several genes organized as a supergene with an hemizygous genetic architecture (Li et al. 2016, Shore et al. 2019, Gutiérrez-Valencia et al. 2022); and ii) species with homomorphic SI systems typically containing large numbers of reproductive groups that are morphologically indistinguishable, with over twenty to hundreds of distinct SI specificities commonly found in natural populations (Lawrence 2000). In the latter scenario, male and female recognition specificities have been found to be encoded by tightly linked but distinct genes, the molecular function of which are highly diverse among the studied SI systems (Takayama & Isogai 2005). The high level of multiallelism in homomorphic SI systems is a theoretical expectation arising from the strong negative frequency-dependent selection acting on these genetic systems (Wright 1939). In contrast, in heteromorphic SI systems, a low number of groups is expected due to the binary nature of the hemizygous genetic architecture, and the phenotypic optimization of the deposition of pollen along the anterior and posterior parts of the pollinators body (Darwin 1877, Barrett and Shore 2008).

Despite these two well-established genetic architectures, a striking evolutionary conundrum was recently spotted in members of the Oleaceae family, with the discovery of a highly unusual SI system, comprising only two categories of mating partners, but in the absence of any discernible morphological differences between the two categories (called Ha and Hb). This homomorphic but diallelic SI (DSI) system has remained stable for a long evolutionary time, since trans-specific pollination assays demonstrated that the two compatibility groups are shared at the phenotypic level among genera as distant as *Phillyrea* (Saumitou-Laprade et al. 2010), *Olea* (the olive tree, Saumitou-Laprade et al. 2017), *Fraxinus* (Vernet et al. 2016) and *Ligustrum* (De Cauwer et al. 2022), representing from 20-30 Myrs (Unver et al. 2017) to 50 Myrs (Olofsson et al. 2019) of divergence. Genetic mapping in *P. angustifolia* (Carré et al. 2021) and in *O. europeae* (Mariotti et al. 2020) showed that the DSI determinants segregate as a single mendelian locus at orthologous positions, but the genomic and molecular bases of this highly unusual SI system have remained elusive.

In this study, we obtained chromosome-scale genome assemblies of *P. angustifolia* individuals belonging to the two SI groups. We identified a 500-700kb hemizygous region also segregating in the olive tree and containing a single conserved candidate gene. This gene is predicted to interfere with the gibberellin pathway and its presence/absence shows perfect association with SI phenotypes across six distantly related genera within the Oleae tribe. Together, our results identified a key determinant of the DSI system, which is largely conserved across this economically important family of plants and has the unique feature of controlling both male and female specificities.

## RESULTS

### The S-locus in *P. angustifolia* corresponds to a 543-kb segregating indel

The S-locus is structurally complex in most species where it has been investigated (*e.g.* Goubet et al. 2012 in Arabidopsis, Wu et al. 2020 in Petunia), so we generated and annotated high-quality chromosome-scale genome assemblies for two selected *P. angustifolia* individuals that are homozygous at the SI locus (S-locus), with S1S1 and S2S2 genotypes, respectively (Billiard et al. 2015). Note that the existence of homozygous S2S2 genotypes is rendered possible by the universal compatibility of males in this species (Saumitou-Laprade et al. 2010). We produced a reference-level genome assembly for the first individual by combining HiFi PACBIO sequences and optical mapping datasets to obtain a hybrid scaffolding assembly, which we organized into 23 pseudo-chromosomes using the *P. angustifolia* genetic map of Carré et al. (2021, Figure S1). We obtained and assembled PACBIO HiFi reads from the second individual (S2S2), and organized the resulting contigs by comparison to the reference. Given the high heterozygosity in the species (Figure S2), the assembler produced two alternative assemblies for each individual (referred to as “hap1” and “hap2”) containing the long and short “allelic” versions of the sequences at each node of the assembly graph.

The genetic map of Carré et al. (2021) predicted that the S-locus was located at position 53.619 cM on linkage group 18, along with seven fully linked Genotyping By Sequencing (GBS) markers. We used blast to position these GBS markers, as well as the flanking recombinant markers, on the *P. angustifolia* genome. This allowed us to delimit the S-locus to a 5.45Mb genomic interval between 12,480,240 - 17,934,613bp on chromosome 18 (Figure 1A). This region corresponds to a chromosomal interval with low recombination (Figure 1A), low gene density and high TE content (Fig 1B), probably corresponding to a centromeric region. The interval contains 65 predicted protein-coding genes, and we hypothesized that it contains the SI determinants.

**Figure 1.**
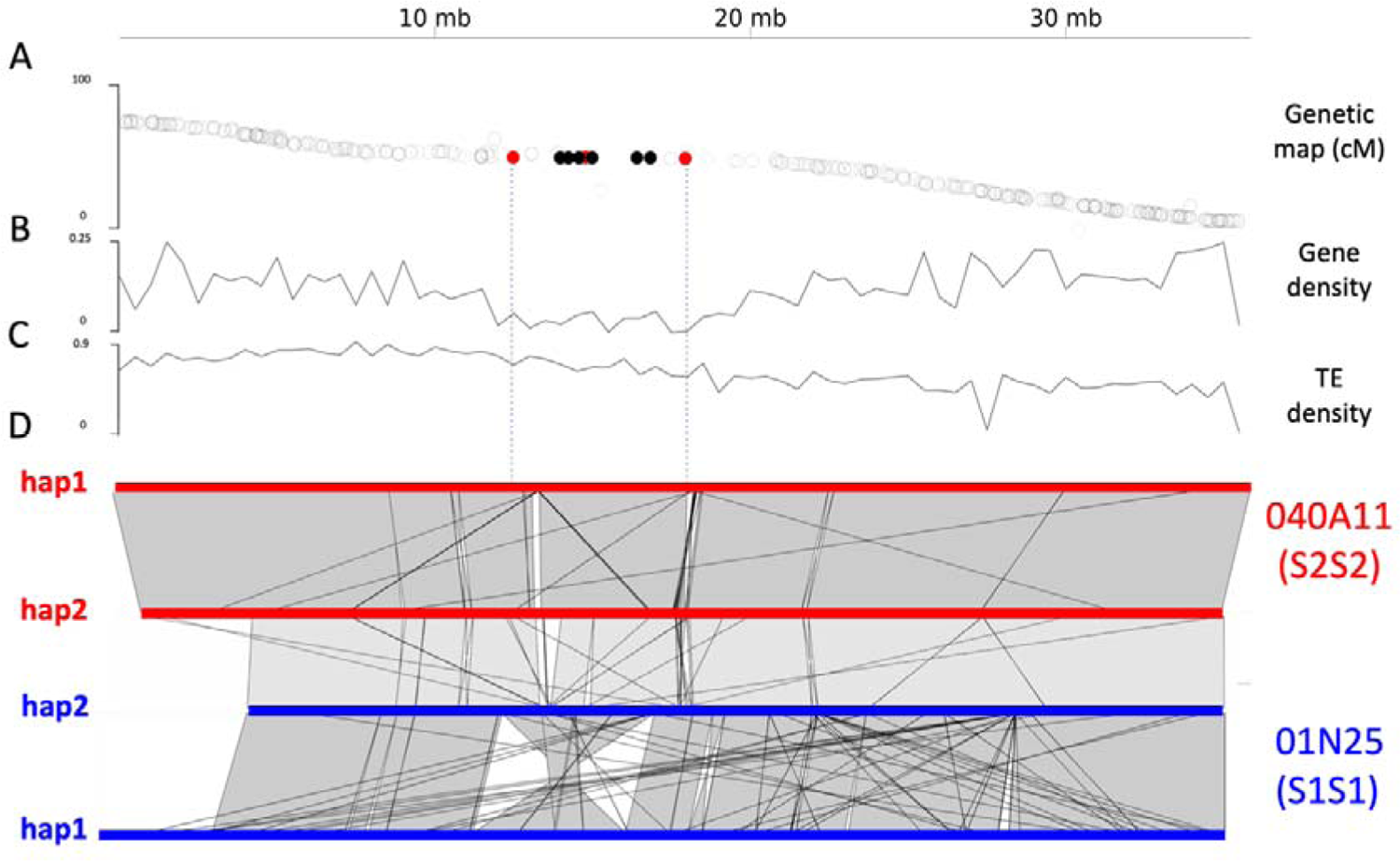
The S-locus maps to a 5Mb centromeric interval. **A.** Comparison of the genetic-to-physical map (in the Y and X axes, respectively) along chromosome 18. GBS markers fully linked to the S-locus (genetic map location: 53.619 cM) are represented in black; those within 0.25cM to the S-locus are represented in red, the others are represented as open black circles. The dotted vertical lines indicate the physical interval between the markers within 0.25cM of the S-locus. **B.** Density of predicted protein-coding genes along chromosome 18. **C**. Density of annotated transposable elements along chromosome 18. **D.** Full chromosomal alignment between the two haplotypes (hap1 and hap2) in each of the two assemblies, indicating that only one of the two S1 chromosomes carries an inverted S-locus. The alignment between hap2 haplotypes from the two genomes is represented by the lighter gray area.

We first noted the presence of an inversion of about 5Mb in the primary assembly of the S1S1 individual, approximately corresponding to the S-locus region (Figure 1C). To evaluate the possibility that the inversion itself could be related to the SI determinant, we computed the rate of synonymous divergence (*K*_S_) between the protein-coding genes in the S1 and S2 haplotypes along chromosome 18. We could not detect any elevation of *K*_S_ for genes inside the inversion as compared to the rest of the chromosome (Figure S3), suggesting that the inversion is indeed very recent and probably unrelated to SI determination given the ancestrality of the two allelic specificities (Vernet et al. 2016). In fact, comparing the two S1 haplotypes of the S1S1 assembly revealed that only one of these (hap1) contained the reverse orientation (Figure 1D). The second haplotype (hap2) had the same orientation as that in the S2S2 assembly, where this “standard” orientation was homozygous. Re-mapping of raw HiFi PACBIO and BioNano signals onto the assembly of the S1S1 individual, and inspection of the inversion breakpoints confirmed that the inversion was truly heterozygous in the sequenced individual, and did not correspond to an assembly error (Figure S4). Finally, to further investigate the presence of this inversion, we produced Nanopore reads from high molecular weight (HMW) genomic DNA of a third *P. angustifolia* individual from the same population (Fabrègues) with a S1S2 genotype. By mapping the raw unassembled reads on the breakpoints of the inversion, we confirmed that the inversion was absent from the S1S2 individual, and is therefore specific to only one of the two S1 chromosomes of the sequenced S1S1 individual (Figure S4). Hence, this inversion appears to be very recent, since it is not fixed among S1 chromosomes, and thus should be independent from the SI determinants that presumably have a single origin, and have remained remarkably stable over extended evolutionary times (Vernet et al. 2016).

To precisely delimitate the region containing the SI determinants (Figure 2A), we used polymorphism revealed by RNA-seq data collected from entire flower buds and pistils, considered as biological replicates. We tested whether any of the 65 predicted protein-coding genes located in the previously identified 5.45Mb genomic interval contained SNPs strictly associated with given SI phenotypes, using fourteen *P. angustifolia* individuals from the same local population (Fabrègues). Within these fourteen individuals, eight were phenotyped by controlled pollination assays as S1S1 (Ha) and six as S1S2 (Hb) (Table S4). We could not find a single fully associated SNP in the whole interval, hence excluding each of these protein-coding genes as the SI determinants.

**Figure 2.**
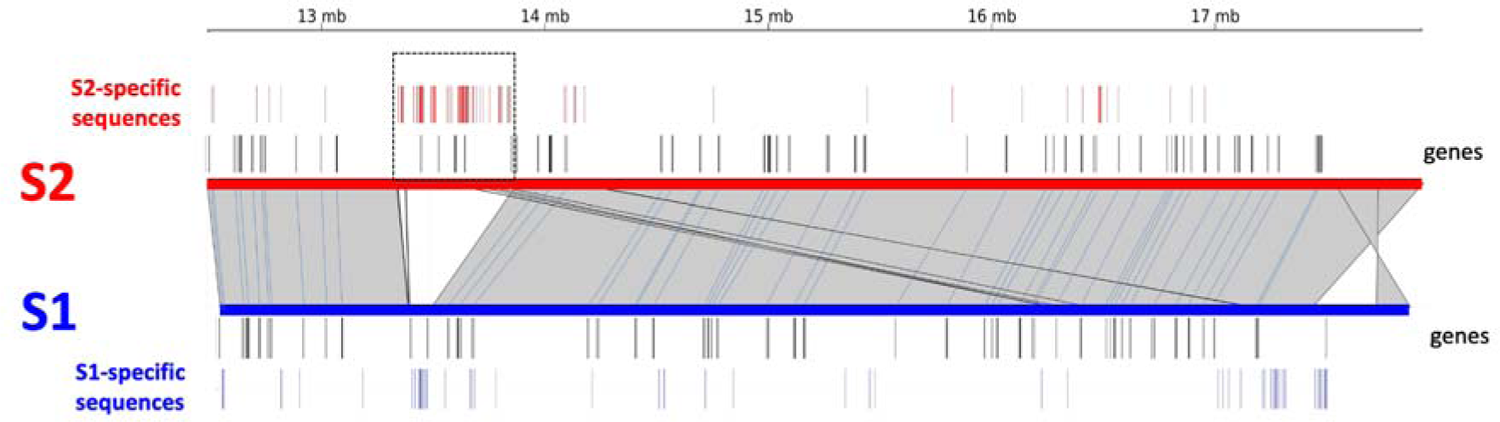
The S2 chromosomes carry a 543kb indel. The candidate 5Mb interval contains 65 predicted protein-coding genes, and a 543kb indel (box delimited by dotted line) spanning over six genes contains a large number of S2-specific sequences. Hap1 of the S2S2 individual was aligned on hap2 (non-inverted) of the S1S1 individual. Gray areas delimited by thin black lines connect aligned portions of the two chromosomes. Predicted protein-coding genes are represented by black vertical lines along each of the two chromosomes. The positions of the complete set of S1- and S2-specific 300bp sequences are represented by blue and red vertical lines, respectively.

Finally, we performed a systematic identification of S1- and S2-specific genomic sequences in the chromosomal interval by blasting for sequence fragments (full set of all non-overlaping 300bp fragments) that were specific to each of the two assemblies (either locally in the S locus genomic interval, or elsewhere across the genome). We found an overall high local sequence similarity between the two assemblies, with the notable exception of a 543kb indel containing an accumulation of sequences specific to the S2S2 assembly (Figure 2), suggesting a hemizygous genetic architecture for the S-locus of *P. angustifolia*. This 543-kb indel contains six predicted protein-coding genes.

### The S-locus hemizygous indel is shared with Olea, and contains a single conserved gene

In order to test for a shared genetic architecture of the DSI system, we then compared the region identified in *P. angustifolia* with the orthologous region in the recently published olive tree genome, *O. europaea, var Arbequina* (Rao et al. 2021). This cultivar is known to belong to SI group Hb (referred to as G1 in Saumitou-Laprade et al. 2017) and thus to have the S1S2 genotype at the S-locus. We found overall strong conservation of the gene content and order along chromosome 18 between the two genomes, but poor conservation of intergenic regions (Figure 3), as expected after *ca.* 32 Myrs of divergence between Olea and Phillyrea (Olofsson et al. 2019). To determine if a segregating indel was also present at the orthologous position in Olea, we remapped short reads from four S1S1 and four S1S2 olive tree cultivars on the Arbequina reference (Jimenez-Ruiz et al. 2020, Table S3). We observed essentially no mapping of short reads from the S1S1 cultivars over a 756-kb fragment, while short reads from the S1S2 cultivars showed a consistent mapping density of about half that of the chromosomal median (Figure 3). Hence, it appears that in the olive tree, the SI groups are also associated with a hemizygous indel at an orthologous position to that of *P. angustifolia*. Interestingly, the relative sizes of the indel in Phillyrea and in Olea (543 vs. 756kb, *i.e.* a ratio of 0.72) are largely consistent with their overall genome size differences (803Mb *vs.* 1.3Gb, *i.e.* a ratio of 0.62, Rao et al. 2021).

**Figure 3.**
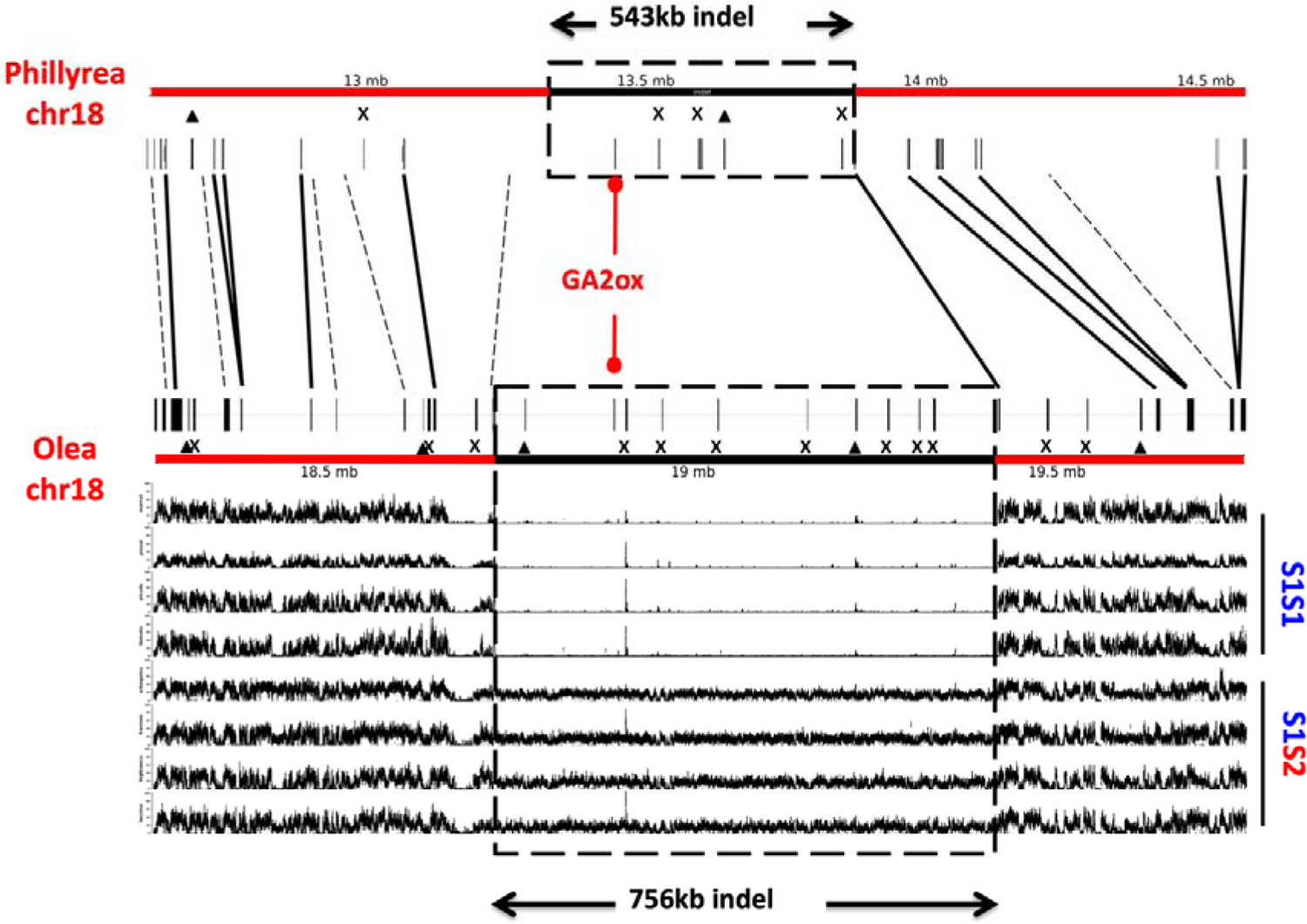
The indel in *P. angustifolia* contains six predicted protein-coding genes. Comparison of the indel sequence between *P. angustifolia* and *O. europeae* (var. Arbequina) reveals that GA2ox is the only conserved gene in the indel, with high divergence of intergenic sequences. Short reads mapping of *O. europaea* accessions identifies a segregating 756kb indel. Triangles indicate annotated genes that putatively correspond to transposable elements. Crosses indicate genes with no sequence similarity (by blast) with anything in the chromosomal fragment of the other species. Solid lines (black and red) indicate orthologous genes. Interrupted lines indicate genes in one species with strong sequence similarity but no gene annotation in the chromosomal fragment of the other species.

Given that *P. angustifolia* pollen can trigger a robust and allele-specific SI response in Olea and *vice-versa* (Saumitou-Laprade et al. 2017), we then reasoned that the molecular determinants of SI should be conserved among these two species. Therefore, we compared the gene content of the indels of Phillyrea and Olea. Strikingly, we found that among the six annotated genes in the Phillyrea indel and the thirteen annotated genes in the Olea indel, only one was common between the two species: a gibberellin-2-oxydase gene (*GA2ox,*), according to the *O. europeae* annotation (Figure 3), and referred to as *G2BD-S* in the companion paper by Raimondeau et al. Analysis of the RNA-seq data from the fourteen *P. angustifolia* individuals of Fabrègues revealed that the *P. angustifolia GA2ox* (*PaGA2ox*) was also the only one of the six genes present in the indel to show consistent expression in buds and pistils that is specific to all six S1S2 individuals. Hence, stringent evolutionary filtering has resulted in the preservation of a single protein-coding gene. Considering that the SI functional response is conserved between the two species, these observations establish *GA2ox* as a prime candidate for the control of SI.

### The presence/absence polymorphism of the *GA2ox* gene is stably associated with SI groups across Oleaceae

To expand the phylogenetic scale of our analysis, we retrieved published genome assemblies from twenty Oleaceae species belonging to five genera and available from NCBI (Table S2). We used BLAST to recover the sequence of the *PaGA2ox* orthologs in the other species. We found that *GA2ox* was present in exactly half of the assemblies as expected (ten out of twenty, Table S2), given that both SI groups (S1S1 and S1S2) are predicted to reach population frequencies of one half in each species. The phylogeny of the sequences obtained covers four genera of the Oleeae tribe (Phillyrea, Olea, Fraxinus, Syringa) and strictly follows the species phylogeny (Figure 4A).

**Figure 4.**
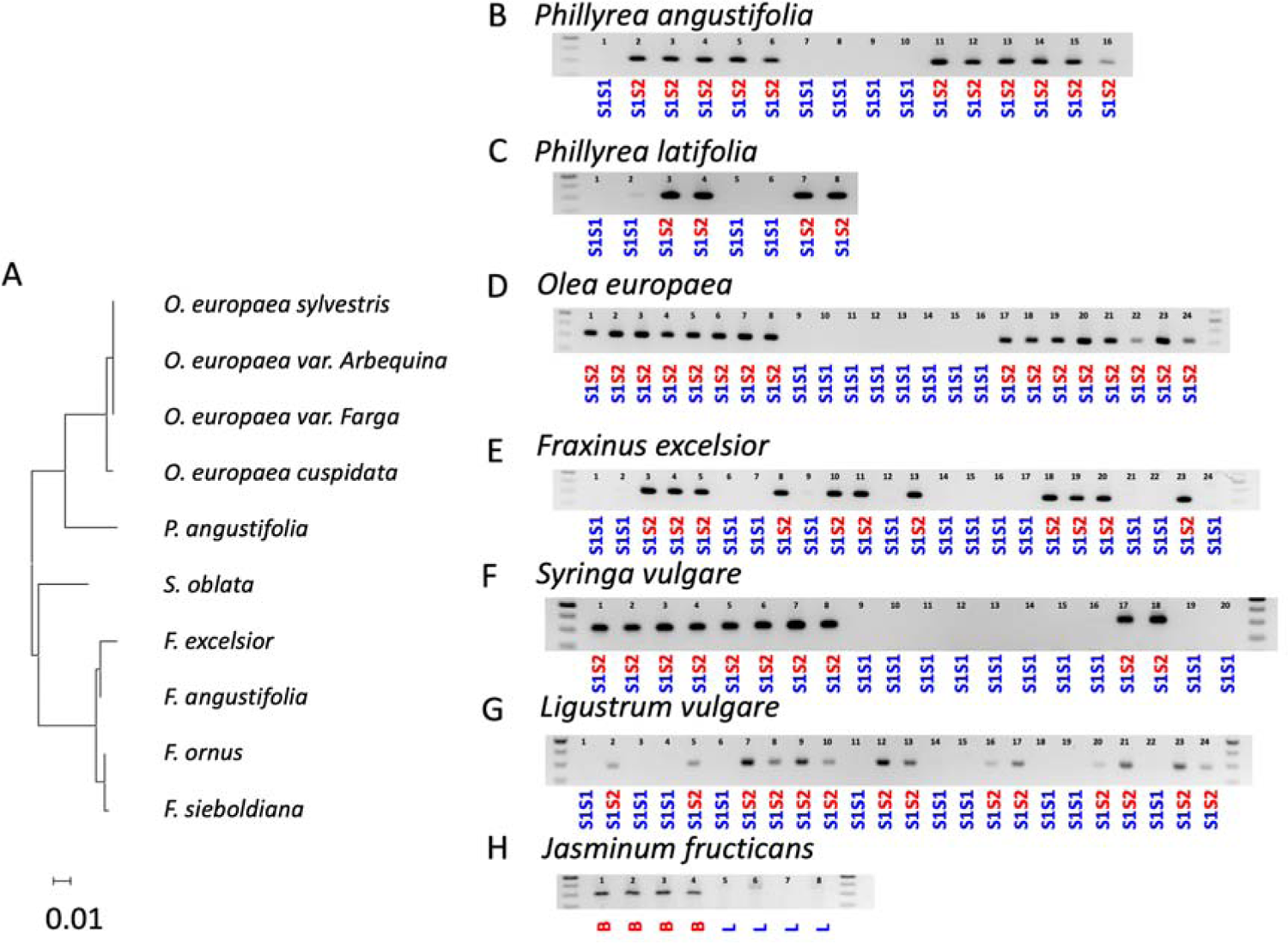
Presence of GA2ox is stably associated with SI phenotypes across distant oleaceae species. **A.** Maximum likelihood phylogeny of the first exon of GA2ox across ten oleaceae species. Presence / absence of PCR products fully correlate with SI groups **B.** in *P. angustifolia* from distant populations. C. in a distinct *Phillyrea* subspecies (*P. latifolia*) D. in a diverse set of *O. europaea* accessions E. in *Fraxinus excelsior.* F. in *Syringa vulgaris*. G. in *Ligustrum vulgare.* H. in *Jasminum fructicans* (B: brevistylous, L: longistylous). The full list of samples tested is reported in Table S6. Numbers above the sample lanes are provided for cross-referencing with Table S6.

Definitive proof that *GA2ox* is the SI determinant requires demonstrating its association with SI groups in divergent species, despite the millions of years of opportunities for recombination. To achieve this, we used the *GA2ox* sequences obtained above to design highly specific PCR primers and tracked the presence of *GA2ox* in samples for which we had previously determined the SI groups by phenotypic assays (Saumitou-Laprade et al. 2010, Vernet et al. 2016, Saumitou-laprade et al. 2017 Saumitou-Laprade et al. 2018, De Cauwer et al. 2020) or that we newly phenotyped (in *Olea europaea, Syringa vulgaris* and *Ligustrum vulgare*, Table S6). The PCR amplification showed a clear signal that was fully associated with S1S2 genotypes across samples from all six species tested, including 47 *P. angustifolia* accessions from distant populations, 10 *P. latifolia* accessions (a different subspecies of Phillyreae), 132 cultivated *O. europaea* varieties, 38 wild *O. europaea europaea* var. *sylvestris* as well as 26, 36, and 24 accessions of the even more distant *Fraxinus excelsior*, *Syringa vulgaris*, and *Ligustrum vulgare* (Figure 4B, C, D, E, F, and G; Table S6). Notably, linkage was even maintained in the basal and distylous *Jasminum fruticans*, where *GA2ox* was present in all five brevistylous but absent from all seven longistylous individuals that we tested (Figure 4H). This suggests that the former correspond to the S1S2 genotype and the latter to the S1S1 genotype, thus formally establishing a link between the determinants of the DSI and those of distyly.

### The GA2ox gene encodes a class-I gibberellin oxidase expressed in floral buds

Sequence analysis of the S-locus *PaGA2ox* gene indicates that it is a class-I gibberellin 2 oxidase enzyme (Figure S5). GA2ox proteins are specialized in the inactivation of GA precursor molecules at different stages of the GA biosynthesis pathway (Ouelette et al. 2023). Given its sequence similarity with class I GA2 oxidases of other flowering plants, *PaGA2ox* is predicted to degrade “end” products of the GA biosynthesis pathway, including the bioactive forms GA1 and GA4. Two protein domains can be identified in PaGA2ox: the 2-oxoglutarate/Fe(II)-dependent oxygenase domain (2OG-FeII_Oxy), spanning exon 2 and 3, and a DIOX_N domain, specifically located in exon 1. The latter corresponds to the highly conserved N-terminal region of proteins with 2-oxoglutarate/Fe(II)-dependent oxygenase activity. Both these domains are consistently found in GA2 oxidases of other flowering plants and are a defining characteristic of this gene family (Figure S5, Cheng et al. 2021). qPCR assays confirmed that *GA2ox* transcripts could only be detected in S2-carrying individuals (Figure S6), with robust expression in mature pistils. Expression in mature anthers was also detectable in one of the three sampled biological replicates. Due to limitations in the size of immature anthers and pistils, we were not able to dissect these tissues at earlier developmental stages, where SI determination could potentially occur. Nevertheless, these results suggest that *PaGA2ox* has the potential to be expressed in both female and male reproductive tissues. Additionally, and as expected, we could not detect any expression in non-reproductive tissues (leaves) of any individual (Figure S6).

### Treatment with gibberellin disrupts the pollen and stigma SI response in an S-allele specific manner

Overall, our results suggest a simple model under which S1S1 pistils and pollen would have basal levels of GA hormone, leading to the Ha specificity phenotype by default. Based on this model we hypothesize that expression of *GA2ox* in S1S2 individuals should modify the balance between the bioactive and the inactive forms of GA, eventually causing these individuals to display the Hb specificity phenotype. To test this, we applied exogenous GA3 on immature *Ligustrum vulgare* floral buds and examined the resulting pollen and pistil SI specificities when the flowers opened. We chose *L. vulgare* because the structure, morphology and size of its inflorescences allow easy treatment with GA3 by inflorescence dipping. We observed that the GA treatment disrupted the SI reaction in both S1S1 and S1S2 individuals, and rendered them self compatible (Figure 5 panels f and p). However, the organ responsible for the breakdown of SI (pollen *vs.* pistil) differed between the two groups. The pistil specificity of the treated S1S1 floral buds remained intact (they still rejected pollen from untreated S1S1 individuals and allowed germination of pollen from untreated S1S2 individuals, Figure 5e and 5g), while pollen specificity switched entirely as a result of the treatment (now triggering a SI reaction when deposited on untreated S1S2 pistils but germinating successfully on untreated S1S1 pistils, Figure 5j and 5b). Strikingly, we observed the exact reciprocal pattern upon treatment of the S1S2 floral buds, with pollen specificity remaining unchanged upon GA treatment (still rejected on untreated S1S2 pistils and successfully germinating on untreated S1S1 pistils, Figure 5l and 5d), whereas pistil specificity entirely switched as a result of the treatment (now triggering a SI reaction with untreated S1S1 pollen, while allowing successful germination of untreated S1S2 pollen, Figure 5m and 5o). Hence, we conclude that GA treatment has a specific effect on the SI phenotype of both groups, switching the pollen specificity of one group (Ha) but the pistil specificity of the other group (Hb).

**Figure 5.**
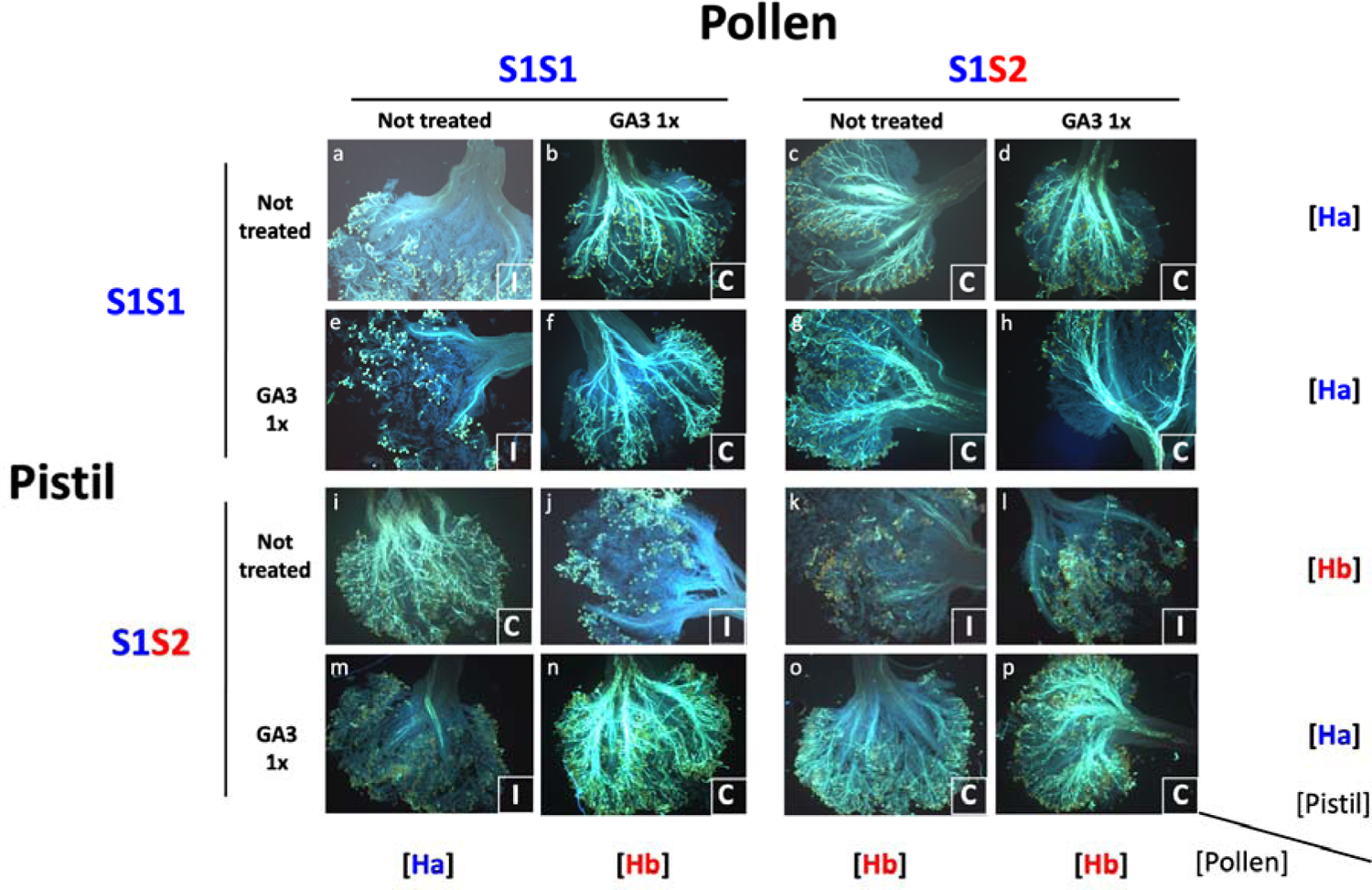
Treatment with gibberellin disrupts the SI response in an S-allele specific manner in *Ligustrum vulgare*

## DISCUSSION

### Comparative genomics suggests a SI system with a single determinant gene

By comparing the nucleotide sequences of the segregating indel in Phillyrea and Olea (separated by 32.22 Myrs), we show that a single gene, *GA2ox*, is conserved, and we find that the presence/absence polymorphism of this gene is stably associated with SI specificities across all species and genera that we tested. Because the SI determinants are shared between Olea and Phillyrea (pollen from one species can trigger the SI response in the other species, and *vice-versa*), these observations provide unambiguous evidence that the *GA2ox* gene acts at the SI determinant.

The conservation of a single gene is striking, as to our knowledge all SI systems so far elucidated at the molecular level entail the action of at least two separate genes, determining the male and female specificities, respectively (Fujii et al. 2016, Rohner et al. 2023). These determinants vary greatly in the molecular functions they encode (Broz and Bedinger 2021) and in their level of repetition, with the male determination being, for example, spread across multiple tandemly duplicated paralogs in Solanaceae (Kubo et al. 2015). Yet, it has never been reported that the male and the female functions are both encoded by a single determinant gene. Although several protein-coding genes are predicted to be present within the S-locus indel in both Phillyrea and Olea, our study clearly shows that only the *GA2ox* gene is conserved. The possibility remains that another kind of genetic element could act as the second determinant, such as a locus producing a non-protein coding RNA. While the action of small non-coding RNAs has been reported at other SI loci, such as at the Brassicaceae S-locus (Tarutani et al. 2011, Durand et al. 2014), they do not directly encode allelic specificities, but rather control their expression at the transcriptional level. In addition, direct sequence comparison of the whole indel nucleotide sequences revealed no other sequence motif that could carry such a conserved element. This interpretation is reinforced by our observation that an exogenous GA3 treatment switched allelic specificities of the male function of S1S1 individuals and of the female function of S1S2 individuals. Hence, at this stage we hypothesize that *GA2ox* acts both as the male and the female determinant of SI specificities, which is a very unusual situation for SI systems. Formal proof of this possibility will ultimately require the obtention of knock-in and knock-out mutants by genetic transformation. While the extended development time of Oleaceae species heavily constrains these experimental approaches, our work provides the solid foundation required for the design of such experiments.

### Mechanistic understanding of how *GA2ox* controls SI specificities is still lacking

So far, only a handful of SI systems have been deciphered at the molecular level (Fujii et al. 2016). In this context, the role demonstrated here for GA represents a fully novel pathway for the control of SI. Interestingly, the implication of a plant hormone is reminiscent of the role of Brassinosteroids, which are involved in the determination of female specificity in Primula (Huu et al. 2016, 2022) and Turnera (Matzke et al. 2021). Nevertheless, it is not yet elucidated how plant hormones can establish SI specificities.

The function of class I GA2 oxidases is usually to modify the balance between the bioactive and inactive forms of gibberellic acids. Their substrate is either bioactive forms of GA, or their immediate precursors, so it is expected that the presence of the *GA2ox* gene in S1S2 individuals lowers the levels of bioactive GA in their reproductive tissues. Even though most players involved in the determination of allelic specificities in Oleaceae remain to be identified, our data shows that the *GA2ox* gene participates in this pathway, likely establishing asymmetries in GA levels, and consequently leading to different specificities between the S1S1 (Ha) and S1S2 (Hb) individuals. This is supported by the fact that changing GA levels in immature floral buds through exogenous supplementation with GA3 triggers changes in female and male specificities: pistils of treated S1S2 (Hb) individuals transition into an Ha phenotype; and pollen of treated S1S1 (Ha) individuals transitions into an Hb phenotype. While it is clear that levels of GA are involved in the determination of allelic specificities, it is surprising that exogenous application of GA changes specificities in a tissue-specific manner: we expected to observe a switch in allelic specificity of both pistil and pollen; however, only one of the two tissues changed specificity upon GA treatment in both S1S1 and S1S2 individuals. This could be explained by tissue-specific dysregulation of gibberellin-related genes as a response to the GA treatment, since GA homeostasis is tightly controlled by several complex feedback mechanisms (Yamaguchi 2008). It is also possible that *GA2ox* would act instead as a dominant negative mutation (Veita 2007), antagonizing the activity of other gibberellin oxidases elsewhere in the genome. Thus, our results call for a more comprehensive exploration of the biochemical activity of the GA2ox protein as well as of the physiological link between the presence of GA2ox, balance between the bioactive and inactive forms of GA in the male and female tissues of both SI groups, and the ultimate fate of compatible *vs.* incompatible pollen. In the meantime, our observation that GA treatment can efficiently compromise the SI reaction holds promise to accelerate breeding programs and optimize olive production in conditions when availability of compatible pollen is limiting.

### Hemizygosity and the long-term maintenance of diallelism

The maintenance of only two S-alleles in the Oleaceae has been described as an evolutionary puzzle, and suggests a long-term evolutionary constraint on the diversification of S-alleles (Billiard et al. 2011, Francq et al. 2023). Strong natural selection in favor of the emergence of new S-alleles is expected to lead to the rapid diversification of S-allele lineages, especially when their number is low (Gervais et al. 2011; Harkness et al. 2021), and SI systems in most plant families typically indeed diversify to a spectacular extent (Lawrence 2000). Why then does this phenomenon not take place in the Oleaceae? A first straightforward explanation lies in our finding that the two S-alleles correspond to the presence or absence of a segregating indel. Such a pattern of presence/absence inherently represents a binary outcome, hence with only two possible states. However, our model posits that the *GA2ox* gene in the indel controls the quantities of bioactive GA, so a continuous variation could in principle be an option. We hypothesize that the determination of SI specificities by GA might work as a bistable switch, rendering more than two alternative states unlikely. Pathways with two distinct and steady stable states can be found in the regulation of cell cycle progression, cell differentiation and phase transitions (Xiong and Ferrell 2003, Ferrell and Machleder 1998, Abley et al. 2021, Topham et al. 2017), and several of these systems integrate hormones as key regulators.

Distyly is another example where two alternative SI specificities are controlled over the long term by a segregating indel, as documented in Primula (Li et al. 2016), Turnera (Shore et al. 2019) and Linum (Gutierrez-Valencia et al 2022). In our study, the stringent evolutionary filtering that occurred over long evolutionary times provided a powerful way to isolate the functional elements necessary for SI determination. In contrast, and in spite of decades of experimental work on distylous plants, the contribution of each of the genes in the indel to the male and female SI specificities *vs.* to the various morphological differences between longistylous and brevistylous individuals remains incompletely elucidated (Kappel et al. 2017, Huu et al. 2022). Interestingly, basal Oleaceae species also exhibit distyly, and our analysis suggests that presence of the indel is associated with the two morphs in Jasminum (see also the companion paper by Raimondeau et al. 2023). In this context, it is possible that the DSI represents a degenerate distyly supergene having lost the morphological determinants of the floral polymorphism. Alternatively, the homomorphic DSI may represent the ancestral condition at the basis of the Oleaceae, with reciprocal differences in style and anther lengths having evolved secondarily several times independently in the basal species. Exploration of the S-locus sequence in distylous species will be a fascinating perspective to determine whether the Olea or Phillyrea indel sequences contain remnants of the genes controlling morphological differentiation. Finally, we note that in heterostylous genera, the brevistylous morph has been identified as associated with the dominant S-locus haplotype in almost all cases investigated (Ganders 1979). Accordingly, just like in Primula and Linum, we found that the presence of the indel in the heterostylous Jasminum seems to be associated with the brevistylous morph. Once we better understand the developmental underpinning of the presence/absence polymorphism in these different systems, it will be interesting to determine whether this is more than just a coincidence.

### DSI and the diversity of mating systems in Oleaceae

A fascinating feature of the Oleaceae is the diversity of their mating systems (Francq et al. 2023), ranging from no sexual differentiation (pure hermaphroditism) to full separation of sexual functions (dioecy), and comprising multiple instances of mating systems that are extremely rare in the rest of the living world, such as androdioecy (Charlesworth 1984, Pannell 2002, Wallander 2008). Here we focused on the DSI determinants, but our results open the way to more detailed studies on the relationship between DSI and mating systems (Francq et al. 2023). An interesting case study of such interactions will be the universal compatibility of pollen produced by male individuals in *P. angustifolia*, which has been shown to be key to their evolutionary success (Saumitou-Laprade et al. 2010). The molecular control of SI based on GA levels uncovered here provides an ideal foundation for the study of the molecular determinants of this important, but mysterious, epistatic interaction. More generally, in some Oleaceae species the determinants of sexual differentiation are independent from the S-locus (*e.g.* in *P. angustifolia*, Carré et al. 2021), while in others they appear to be fully linked (*e.g.* in *F. excelsior*, Saumitou-Laprade et al. 2018). Understanding the consequences of the different evolutionary trajectories followed by these species of such an economically important plant family will now be essential.

## Acknowledgements

Sequencing of the *P. angustifolia* genomes was funded by France_Olive as part of the study agreement entitled “Study of the self-incompatibility locus in olive and Phillyrea” to PSL. Resources for this work were also partly provided from the European Research Council (NOVEL project, grant #648321) to VC and ANR TE-MoMa (grant ANR-18-CE02-0020-01) to VC and SL, as well as an EMBO fellowship (ALTF 657-2020) to RAB. We would like to thank Antònia Nino from the IRTA institute, Hélène Lasserre and Julien Balajas from France Olive for collecting inflorescences and providing access to the olive cultivars and oleaster collections. We would like to thank Arnaud Dowkiw and Lemonniers Nurseries for access to *Fraxinus excelsior* samples, Hanno Schaefer for collecting *Ligustrum vulgare* inflorescences in the Isar Valley (Germany), Amelia Dumbravă for access to *Syringa vulgaris* populations in the Portes de Fer Natural Park (Romania). Finally, we would like to thank Michel and Christine Prat for their warm welcome and for collecting *Jasminum fruticans*. The authors thank Luciana Baldoni for her contribution to the study design at the early stages of the project, as well as Jacques Lepart^†^, Mathilde Dufay and Patrick Achard for scientific discussions. This work was performed using the infrastructure and technical support of the “Plateforme Serre, cultures et terrains expérimentaux – Université de Lille” for the greenhouse/field facilities. We thank David Degueldre and the staff of the “Plateforme des Terrains d’Expérience du LabEx CeMEB” (CEFE, CNRS) for plant cultivation.

## Author contributions

Conceptualization: PSL and PV; Methodology: SL, RAB, XV; Investigation: PSL, VC, RAB, AC, AT, WM, SS, CG, SM, RM, CM, SL; Software: CM, SG, AR; Resources: AB; Writing – Original Draft: VC, PSL, AC, RAB, AT, WM, SM, RM, AR; Writing – Review & Editing: VC, PSL, SB, XV, PV, RAB; Funding Acquisition: PSL, VC; Supervision: PSL

## METHODS DETAILS

### Biological material and SI genotype determination

We generated high-quality chromosome-scale genome assemblies of two *P. angustifolia* individuals. The first one (01N25 in Billiard et al. 2015) was a hermaphrodite plant belonging to SI group Ha (Billiard et al. 2015), with S-locus genotype S1S1, as determined by the segregation of SI groups in its progeny. The other one (040A11 in Billiard et al. 2015) was a male individual with genotype S2S2, also as determined by the segregation of SI groups in its progeny. A third individual (13A06 in Carre et al. 2021) was sequenced using the Oxford Nanopore Technology. This plant is a male, and was used as pollen donor to generate the controlled progeny to produce the GBS map in Carre et al (2021). According to the segregation of SI groups in its progeny, this plant has a heterozygous genotype (S1S2) at the S-locus. The ancestors of these individuals (parents or great parents) originate from a local population in Fabrègues (southern France).

### HMW DNA isolation

High molecular weight (HMW) DNA was extracted from frozen leaves using QIAGEN Genomic-tips 500/G kit (Qiagen, MD, USA), following the tissue protocol extraction. Briefly, 2g of young leaves material were grounded in liquid nitrogen with mortar and pestle. After 3h of lysis and one centrifugation step, the DNA was immobilized on a column. After several washing steps, DNA was eluted from the column, then desalted and concentrated by isopropyl alcohol precipitation. A final wash in 70% ethanol was performed and the DNA was resuspended in EB buffer. DNA quantity and quality were assessed using NanoDrop and Qubit (Thermo Fisher Scientific, MA, USA). DNA integrity was also assessed using the Agilent FP-1002 Genomic DNA 165 kb on the Femto Pulse system (Agilent, CA, USA).

### PACBIO sequencing and assembly

High Fidelity (HiFi) libraries were constructed using the SMRTbell® Template Prep kit 2.0 (Pacific Biosciences, Menlo Park, CA, USA) according to PacBio recommendations (SMRTbell® express template prep kit 2.0 - PN: 100-938-900). HMW DNA samples were first purified with 1X Agencourt AMPure XP beads (Beckman Coulter, Inc, CA USA), and sheared with Megaruptor 3 (Diagenode, Liège, BELGIUM) at an average size of 20 kb. After end repair, A-tailing and ligation of SMRTbell adapter, the library was size-selected on the BluePippin System (Sage Science, MA,USA) at range sizes of 10-50kb. The size and concentration of libraries were assessed using the Agilent FP-1002 Genomic DNA 165 kb on the Femto Pulse system and the Qubit dsDNA HS reagents Assay kit.

Sequencing primer v5 and Sequel® II DNA Polymerase 2.2 were annealed and bound, respectively, to the SMRTbell libraries. Each library was loaded on two SMRTcell 8M at an on-plate concentration of 90pM. Sequencing was performed on the Sequel® II system at the Gentyane Genomic Platform (INRAE Clermont-Ferrand, France) with Sequel® II Sequencing kit 3.0, a run movie time of 30 hours with an Adaptive Loading target (P1 + P2) at 0.75. HiFi reads were produced with the PacBio Sequel II system on four SMRTCells and were assembled using the HiFiasm assembler (v0.15.5, Cheng *et al*., 2021; https://github.com/chhylp123/hifiasm).

To assess the completeness and quality of the assemblies, we used the Benchmarking Universal Single-Copy Orthologs (BUSCO) pipeline with the viridiplantae database (Simão et al, 2015). We obtained a 99.3% complete BUSCO score on the primary assembly. In addition, we performed k-mer analysis to quality control the dataset using Jellyfish tool (Marcais *et al*, 2011) and the assemblies using module “comp” of the k-mer Analysis Toolkit (Mapleson *et al*., 2017). All the metrics are reported in Table S1.

### Optical map

To achieve a reference-level genome assembly for the first individual (S1S1), we combined the 40X HiFi PACBIO sequences obtained above with 588X optical mapping datasets. Briefly, ultra HMW DNA (uHWM DNA) was purified from 1g of fresh dark treated very young leaves according to the Bionano Prep Plant Tissue DNA Isolation Base Protocol (30068 - Bionano Genomics) with the following specifications and modifications. Briefly, the leaves were fixed in a buffer containing formaldehyde. After three washes, leaves were cut in 2 mm pieces and disrupted with a rotor stator in homogenization buffer containing spermine, spermidine and beta-mercaptoethanol. Nuclei were washed, purified using a density gradient and then embedded in agarose plugs. After overnight proteinase K digestion (Qiagen) in the presence of lysis buffer and a one hour treatment with RNAse A (Qiagen), plugs were washed and solubilized with 2 µL of 0.5 U/µL AGARase enzyme (ThermoFisher Scientific). A dialysis step was performed in TE Buffer (ThermoFisher Scientific) to purify DNA from remaining residues. The DNA samples were quantified by using the Qubit dsDNA BR Assay (Invitrogen). The presence of megabase-sized DNA molecules was visualized by pulsed field gel electrophoresis (PFGE). Labelling and staining of the uHMW DNA were performed according to the Direct Label and Stain (DLS) protocol (30206 - Bionano Genomics). Briefly, labelling was performed by incubating 750 ng genomic DNA with 1× DLE-1 Enzyme for 2 hours in the presence of 1× DL-Green and 1× DLE-1 Buffer. Following proteinase K digestion and DL-Green clean-up, the DNA backbone was stained by mixing the labelled DNA with DNA Stain solution in the presence of 1× Flow Buffer and 1× DTT, and incubating overnight at room temperature. The DLS DNA concentration was measured with the Qubit dsDNA HS Assay (Invitrogen, Carlsbad, CA, USA). Labelled and stained DNA was loaded on 1 Saphyr chip and was run on the BNG Saphyr System according to the Saphyr System User Guide. Digitalized labelled DNA molecules were assembled to optical maps using the BNG Access software (solve version 3.5). The molecule N50 was 250kb.

### Scaffolding contigs with the optical and genetic maps

A hybrid scaffolding was then performed between the sequence assembly and the optical genome map with the hybridScaffold pipeline (https://bionano.com/wp-content/uploads/2023/01/30073-Bionano-Solve-Theory-of-Operation-Hybrid-Scaffold.pdf; solve version 3.6).

Finally, the genetic map of Carré, et al. (2021) was used to finalize scaffolding. This map consists of an overall total of 15,814 SNPs contained in 10,388 GBS fragments (some GBS fragments contained more than one SNP) genotyped in 196 offspring. We used BLAST (Camacho et al. 2009) to align the sequence of these GBS fragments onto the contigs obtained from the assemblies. These alignments were used to organize contigs containing fragments belonging to the same linkage group and achieve chromosome-scale assemblies. The position of these alignments along the pseudo-chromosomes were then displayed using GViz (Hahne & Ivanek 2016). The SI phenotype (Ha *vs.* Hb) was mapped at position 53.619 cM on linkage group 18 by Carré et al. (2021), and we delimited the chromosomal interval containing the S-locus based on the immediately flanking upstream and downstream markers mapped at position 53.126cM (one GBS sequence, one SNP) and 53.864cM (two GBS sequences, three SNPs), respectively.

### HMW DNA isolation and Oxford Nanopore sequencing

HMW DNA was extracted from leaves of a third *P. angustifolia* individual (13A06), whose S-locus genotype was S1S2. HMW DNA was extracted using the Carlson lysis buffer followed by purification using the QIAGEN Genomic-tip 500/G (Qiagen, MD, USA) with slight modification. 1g of fresh leaves was grinded in a mortar to a fine powder in presence of liquid nitrogen. The powdered material was dispensed into two 50 mL centrifuge tubes containing 20 mL of pre-warmed (65°C) lysis buffer. After one hour of lysis, 20 mL of chloroform has been added to each tube, followed by vortexing and centrifugation. The supernatant was collected, mixed with 0.7x volumes of isopropanol and pelleted by centrifugation. The pellets were resuspended with 19 mL of G2 buffer, from the QIAGEN Blood and Cell Culture DNA Maxi Kit t (Qiagen, MD, USA). Purification was then performed using QIAGEN Genomic-tips 500/G based on the indications of the protocol. Then DNA was precipitated by isopropanol, washed in 70% ethanol and re-suspended in TB buffer. To enhance recovery of long DNA fragments, 9 µg of DNA were processed using the Short Read Eliminator Kit XL (Circulomics, Baltimore, MD, Cat #SS-100-101-01) according to the supplier’s instructions. DNA quantity and quality were assessed using NanoDrop and Qubit (Thermo Fisher Scientific, MA, USA) before and after size selection.

DNA libraries were prepared using the Oxford Nanopore Technologies kit SQK-LSK110 with the following modifications. DNA repair and end-prep (New England BioLabs, Ipswich, MA, Cat #E7546 and Cat #M6630) were performed with 3 µg DNA, in two separate tubes (1.5 µg in each tube) in a total reaction volume of 60 µl each, incubated at 20°C for 60-90 minutes, and 60°C for 60-90 minutes. The two DNA repair and end-prep reactions were combined and cleaned with 120 µl of Agencourt AMPure XP beads (Beckman Coulter, Brea, CA, Cat #A63880) with an incubation time ranging from ten to 20 minutes and an elution time of five to ten minutes. Ligation was performed at room temperature for two hours. The ligation reaction was cleaned using Agencourt AMPure XP beads with an incubation time of ten to 20 minutes and an elution time of 15 to 25 minutes at room temperature or 37°C. Sequencing was performed with the Oxford Nanopore Technologies (Oxford, UK) MinION (MIN-101B) device with a total of 13 FLO-MIN106 Rev D flow cells. The MinKNOW software version 4.3.12 (https://community.nanoporetech.com/downloads) was used to collect data. The running parameters were set to default and the fast basecalling model was used to generate real-time run statistics. After each run, a new basecalling was performed by using Guppy v5.0.13 with the super accurate configuration model on an Intel core I9 workstation equipped with an Nvidia RTX 2080ti GPU. Adapters were trimmed out with Porechop software (https://github.com/rrwick/Porechop).

### Preparation of RNA samples

We performed two replicate RNA-seq experiments on 14 genotypes (8 S1S1 and 6 S1S2) from the Fabrègues population. For each of these individuals, the first experiment was based on five mixed-stages whole flower buds, and the second was based on ten dissected pistils that had been pollinated *in vitro* using pollen from a Hb individual. Samples were ground in liquid nitrogen and total cellular RNA was extracted using a Spectrum Plant Total RNA kit (Sigma, Inc., USA) with a DNAse treatment. RNA concentration was first measured using a NanoDrop ND-1000 Spectrophotometer then with the Quant-iT™ RiboGreen® (Invitrogen, USA) protocol on a Tecan Genius spectrofluorimeter. RNA quality was assessed by running 1 μL of each RNA sample on RNA 6000 Pico chip on a Bioanalyzer 2100 (Agilent Technologies, Inc., USA). Samples with an RNA Integrity Number (RIN) value greater than eight were deemed acceptable.

### RNA-seq library construction and sequencing

The TruSeq RNA sample Preparation v2 kit (Illumina Inc., USA) was used according to the manufacturer’s protocol with the following modifications. In brief, poly-A containing mRNA molecules were purified from 1 ug total RNA using poly-T oligo attached magnetic beads. The purified mRNA was fragmented by addition of the fragmentation buffer and was heated at 94°C in a thermocycler for four minutes to yield library fragments of 250-500 bp. First-strand cDNA was synthesized using random primers to eliminate the general bias towards 3’end of the transcripts. Second strand cDNA synthesis, end repair, A-tailing, and adapter ligation was done in accordance with the manufacturer’s protocols. Purified cDNA templates were enriched by 15 cycles of PCR for 10s at 98°C, 30s at 65°C and 30s at 72°C using PE1.0 and PE2.0 primers and with Phusion DNA polymerase (NEB, USA). Each indexed cDNA library was verified and quantified using a DNA 100 Chip on a Bioanalyzer 2100, then pooled in equimolar amounts by sets of ten samples. The final library was quantified by real-time PCR with the KAPA Library Quantification Kit for Illumina Sequencing Platforms (Kapa Biosystems Ltd, SA) adjusted to 10 nM in water and provided to the Get-PlaGe core facility (GenoToul platform, INRA Toulouse, France http://www.genotoul.fr) for sequencing.

Final pooled cDNA libraries were sequenced using the Illumina mRNA-Seq, paired-end protocol on a HiSeq2000 sequencer, for 2 x 100 cycles. Libraries were diluted to 2 nM with NaOH and 2.5 μL transferred into 497.5 μL HT1 to give a final concentration of 10 pM. 120 μL were then transferred into a 200 μL strip tube and placed on ice before loading onto the cBot. Mixed libraries from ten individual indexed libraries were run on a single lane. The flow cell was clustered using TruSeq PE Cluster Kit v3, following the Illumina PE_Amp_Lin_Block_V8.0 protocol. Following the clustering procedure, the flow cell was loaded onto the Illumina HiSeq 2000 instrument. The sequencing chemistry used was v3 (FC-401-3001, TruSeq SBS Kit) with 2×100 cycles, paired-end, indexed protocol. Image analyses and basecalling were performed using the HiSeq Control Software (HCS 1.5.15) and Real-Time Analysis component (RTA 1.13.48).

### A *de novo* transcriptome assembly for *P. angustifolia*

Demultiplexing was performed using CASAVA 1.8.1 (Illumina) to produce paired sequence files containing reads for each sample in Illumina FASTQ format. Reads were cleaned and filtered with CutAdapt (Martin 2011; option: *--overlap=30*) and Prinseq (Schmieder and Edwards 2011; option: *-min_len 80-trim_tail_left 5-trim_tail_right 5 - lc_method entropy - lc_threshold 70*). The *de novo* assembly protocol was based on Evangistella et al. (2017). Following Wang and Gribskov (2017), we first pre-assembled the reads with Trinity v 2.5.1 and Trans-Abyss v.1.5.5 using the default *K-mer*, *i.e. K-mer = 25* for Trinity (min length=200bp) and *K-mer = 32* for Trans-Abyss, (min length=100bp). Contigs of the pre-assemblies with nucleotide sequence identity above 0.98 were merged using the CD-HIT-EST tool (Li et al. 2006) and verified using Transrate v1.0 (Smith-Unna et al. 2016). These individual *de novo* assemblies were then merged again with Trans-Abyss and we used the EvidentialGene tr2aacds pipeline (Gilbert 2019) to reduce this assembly into a first set of non-redundant unitigs. Following Armero et al. (2017), we used BRANCH (Bao et al. 2013) to improve unitig sequences by aligning the RNA sequencing reads onto the unitigs with a modified version of BLAT (Kent 2002). This latter step identifies novel unitigs, extends incomplete unitigs and joins fragmented ones (Bao et al. 2013). Finally, we sequentially used FrameDP v1.2.2 (Gouzy et al. 2009) and the scripts tr2aacds.pl of the EvidentialGene pipeline (Gilbert 2019) and main.pl (Armero et al. 2017) to remove redundant and/or chimeric unitigs based on their translated polypeptide sequences.

### Annotation of protein-coding genes and transposable elements

We used MAKER (Campbell et al. 2014) to predict protein-coding genes on the two primary assemblies. Briefly, MAKER starts from *ab initio* gene prediction by Augustus v. 3.3.3. (Stanke et al. 2006) trained on the Arabidopsis genome and then searches for a series of additional evidences using the set of predicted proteins and CDS from the olive tree genome (Cruz et al. 2016) as well as unitig sequences from the *de novo P. angustifolia* transcriptome described above. We then aligned raw RNA-seq reads from bud and pistil tissues obtained from fourteen *P. angustifolia* individuals with known SI phenotypes (eight Ha individuals and six Hb individuals, assumed to carry S1S1 and S1S2 genotypes, respectively, Table S4) on the genome using the splice-aware aligner HiSat2 (Kim et al. 2015), and used Stringtie (Pertea et al. 2015) to refine the prediction of transcripts using information from the RNA-seq reads split across intron-exon boundaries. We retained Augustus gene models for which at least one additional evidence was present. We used RepeatMasker to identify TEs based on the olive tree genome (Jimenez-Ruiz et al. 2020; http://olivegenome.org/genome_datasets/Olea_europaea.denovo.library.fa.zip), and we eliminated gene predictions overlapping with TE annotations.

### Genome alignment and sequence comparison

To identify major chromosomal rearrangements, we used minimap2 (Li 2021) to align the alternative assembly of each individual to their respective primary assembly (using the - asm5 option), and to align them to one another. The resulting paiwise alignments were displayed using the *pafr* library in R (https://github.com/dwinter/pafr). We retrieved the sequence interval between the GBS markers aligned on the 040A11 hap1 assembly and used minimap2 with default parameters to align it to the 01N25 hap2 assembly, and displayed the alignment using *pafr*.

To identify nucleotide sequences specific to the S1 or S2 chromosomes, we split the chromosomal intervals between the non-recombining GBS markers linked to the S-locus into consecutive stretches of 300bp and blasted them onto the rest of the genome. We retained only those with no hit above 80% identity and concatenated all overlapping fragments.

### Mapping short Illumina reads from *Olea europaea* accessions

To determine whether the indel we identified in P. angustifolia was also segregating in *O. europaea*, we then used bowtie2 (Langmead and Salzberg 2012) to map publically available short Illumina reads from eight *O. europaea* accessions whose SI phenotype had been determined previously (Table S3) on the complete *O. europaea var.* Arbequina genome (Rao et al. 2021; https://bigd.big.ac.cn/gwh/Assembly/10300/show). We used samtools to vizualise and quantify variation of the depth of aligned sequences along chromosome 18 with a quality threshold of Q30.

### Sequence comparison across distant Oleaceae species, specific primer design, PCR protocol and association study across Phillyrea and Olea accessions

We retrieved assembled genomes from 20 Oleacea species available from the literature (Table S2). We used blast with default parameters to search for *PaGA2ox* orthologs. Based on the aligned sequences of the first exon of the GA2ox orthologs, we designed PCR primers and optimized amplification conditions to track the presence of *GA2ox* in a series of samples with known SI phenotype (Table S6).

### Phylogeny of GA2ox proteins

To place PaGa2ox in the phylogenetic tree of GA2 oxidase enzymes, we collected previously published protein sequences of GA2ox enzymes of different flowering plant species for which the enzyme class was described. The phylogenetic tree was inferred by the Neighbor-Joining method, using MEGA11 (Tamura et al. 2021). The percentage of replicate trees in which the associated taxa clustered together in the bootstrap test (500 replicates) are shown next to the branches. This analysis involved a total of 41 amino acid sequences from *Arabidopsis thaliana* (Cheng et al. 2021), Rice (*Oryza sativa*, Han & Zhu 2011), Grapevine (*Vitis vinifera*, Giacomelli et al. 2013), Tomato (*Solanum lycopersicum*, Chen et al. 2016), Peach (*Prunus persica*, Cheng et al. 2021) and Barley (*Hordeum vulgare*, Ouelette et al. 2023). Domains within the PaGA2ox protein were identified by InterPro (Paysan-Lafosse et al. 2023).

### qPCR and functional characterization of the candidate gene

To study the expression of GA2ox in different *P. angustifolia* tissues we manually dissected anthers and stigmas from closed buds one day before anthesis. We also isolated non-dissected immature buds two weeks before anthesis, as well as leaves. All samples were flash-frozen in liquid nitrogen immediately upon collection and RNA was extracted using the NucleoSpin RNA Plus kit (Macherey-Nagel) following the supplier’s instructions. The RevertAid First Strand cDNA Synthesis Kit (Thermo Scientific) was used to synthesize cDNA and qPCR was performed using the iTaq Universal SYBR Green Supermix (BioRad) on a Lightcycler 480 instrument (Roche). Primer sequences are detailed in Table S5. We selected *PaPP2A* (Protein Phosphatase 2A) to be used as a reference gene, based on its previous validation for use in expression analysis in *Olea europaea* (Ray and Johnson 2014). Relative *PaGA2ox* transcript abundance was estimated using the Pfaffl method (Pfaffl 2001).

### GA2 supplementation experiment

In order to test the effect of GA3 on SI specificities, we took advantage of the relatively large and flexible inflorescences of *Ligustrum vulgare*. We selected three S1S1 (Ha) and two S1S2 (Hb) individuals whose SI phenotype had been characterized in De Cauwer et al. (2021) and cultivated them in an insect-proof greenhouse to avoid pollen contamination of the opening buds. On each individual plant, we selected and labeled four inflorescences - one for each of the treatments described below. Each treatment consisted of immersing the complete inflorescence for a few seconds in a Falcon tube containing 50 mL of solution. We applied four treatments (1) a control with no GA3, (2) a 20µM GA3 solution (0.1x), (3) a 200µM GA3 solution (1x) and (4) a 2mM GA3 solution (10x). Each treatment was applied twice a week from the end of April until the stage “white bud” at the end of May. When an inflorescence contained at least three open flowers, it was immediately collected and transferred to the laboratory for the stigma test. Open flowers were eliminated, and the inflorescence containing only closed buds was kept overnight at 20°C under protection against pollen contamination. After 16 hours the newly opened flowers were collected, emasculated by removing the corollae and planted in agar medium. The corollae containing the two non-dehisced anthers were placed under dry air conditions (for two to three hours) until dehiscence. Then, receptive emasculated flowers in agar and the dehisced anthers were used to perform the different crossing schemes presented in Figure 5. Each cross was performed with five technical replicates.

## Data availability

Data from the present study have been deposited on the NCBI database under bioproject #

PRJNA993866, including

– Genome assemblies and annotations
– Corrected PACBIO reads
– RNA-seq reads
– *de novo P. angustifolia* transcriptome assembly
– Nucleotide sequence of the GBS markers

Accession numbers for raw nanopore reads of the S1S2 individual (13A06):

– Bioproject: PRJNA984176 - Biosample: SAMN35742618

Supplementary figures are publicly available on Figshare at: 10.6084/m9.figshare.23710410.

## SUPPLEMENTARY TABLES

(publicly available at 10.6084/m9.figshare.23710410)

**Table S1**. Detailed description on the sequencing data (PACBIO, nanopore sequencing reads, signals of bionano molecules) and assembly metrics.

**Table S2.** References of the twenty genome assemblies used to obtain the position and sequence of exon 1 of the GA2ox gene.

**Table S3.** Accession numbers for Illumina short reads from eight *O. europaea* accessions from Jimenez-Ruiz et al. (2020).

**Table S4.** Parental origin and S-locus genotype of the 14 *P. angustifolia* samples used for RNA-seq and the three samples used for tissue-specific qPCR.

**Table S5.** Sequences of primers used in this study (presence/absence genotyping and qPCR of *PaGA2ox*)

**Table S6.** Origin of samples used to test the correspondence between presence/absence of the *GA2ox* gene and SI phenotypes in *P. angustifolia*, *P. latifolia, O. europea, F. excelsior, S. vulgaris*, *L. vulgaris, J. fruticans*

## SUPPLEMENTARY FIGURES

**Figure S1.** Congruence between the position of the GBS markers on the reference genome assembly (33 super-scaffolds) and along the genetic map (23 linkage groups, Carré et al. 2022).

**Figure S2.** K-mer analysis by JellyFish is typical of a highly heterozygous genome.

**Figure S3.** Variation of synonymous divergence (*K*_S_) for protein-coding genes annotated along chromosome 18. The vertical lines correspond to the border of the inversion identified between the primary assemblies of 01N25 (S1S1) and 040A11 (S2S2).

**Figure S4.** Mapping of long reads from the three sequenced individuals (01N25: PACBIO, 040A11: PACBIO, 13A06; ONT) on the 5’ and 3’ breakpoints of the inversion in the hap1-01N25 reference (IGV snapshots).

**Figure S5.** Phylogeny of GA2ox enzymes. A. Phylogenetic tree of GA2 oxidases of different flowering plant species. Sequences are divided into three classes corresponding to the three subtypes of GA2 oxidase enzymes (Ouelette et al. 2023). PaGA2ox is highlighted in blue. At, *Arabidopsis thaliana*; Os, *Oryza sativa*; Vv, *Vitis vinifera*; Sl, *Solanum lycopersicum*; Pp, *Prunus persica*; Hv, *Hordeum vulgare*. B. PaGA2ox protein domains identified by InterPro. DIOX_N: non-haem dioxygenase N-terminal domain, 2OG-FeII_Oxy: 2-oxoglutarate Fe(II)-dependent oxygenase superfamily domain.

**Figure S6.** Expression of *PaGA2ox* in different plant tissues. *PaGA2ox* expression levels in leaves, floral buds, stigmas and anthers of *Phillyrea angustifolia*. Expression was measured in 3 different Ha (S1S1) and Hb (S1S2) individuals, and normalized to floral buds of Hb individuals. The presented expression levels correspond to the mean of three technical replicates. *PaPP2a* was used as a reference gene.

